# Counterproductive: coinfection of a water flea by a fungus and a microsporidium reduces the reproductive outputs of all parties

**DOI:** 10.1101/2025.10.29.685260

**Authors:** Snir Halle, Aran Sofer, Frida Ben-Ami

## Abstract

Organisms are often coinfected by more than one parasite species. These coinfections can alter the fitness of both host and its coinfecting parasites. Changes in host fitness are usually estimated by comparing parasite virulence under coinfection to the virulence of each parasite under single infection. While many studies focus on host survival as proxies of virulence, virulence can be expressed via reduced host fecundity. Here, we studied the outcome of coinfection of water fleas (*Daphnia magna*) by two microparasitic species, the fungus *Metschnikowia bicuspidata* and the microsporidium *Hamiltosporidium tvaerminnensis*. We found that *Metschnikowia* expressed its virulence mainly via host mortality and *Hamiltosporidium* expressed it mainly via host reproduction. Despite their competitive interaction, indicated by their reduced spore production, both parasites were able to fully express their virulence during coinfection. Host survival was dominated by *Metschnikowia* while clutch-size was dominated by *Hamiltosporidium*. Furthermore, coinfected hosts experienced increased virulence that was manifested only by additional reduction in reproduction (less clutches were released). Our results demonstrate that focusing only on the survival component of host fitness can miss important outcomes of coinfections. Since reduced host fecundity is a common outcome of parasitism, the influence of coinfection on host reproduction deserves more attention.

## Introduction

In nature, organisms are usually exposed to multiple parasite species and coinfections are ubiquitous and well documented [1]. Due to their capacity to modify the outcome of infection for both host and its parasites, coinfections have been suggested to play an important role in parasite evolution [2,3] and epidemiology [4,5]. These modifications are influenced by numerous factors, such as host traits [6,7], parasite traits [8,9], environmental factors [10,11] and interactions between two or more of these major players [12–14]. This complexity serves as an ongoing call to increase our knowledge of coinfections and their subsequent outcomes.

Changes in host and parasite fitness are the focal point of many coinfection studies. From the parasite’s perspective, the presence of additional parasites can modify its fitness by altering infectivity [15–17] and transmission [18–20]. These changes can be positive [14,21,22] or negative [8,18,23], while in other cases a parasite can be indifferent to the presence of another [24–26]. From the host’s perspective, the presence of additional parasite species can alter the level of pathology it experiences (i.e., virulence) compared to infection by one of the participants. The change in virulence can be positive (i.e., virulence will decrease) [8,10], negative (i.e., virulence will increase) [9,19,27] or simply determined by the more virulent coinfecting parasite [6,25,28]. Although many studies have estimated virulence via changes in host survival (i.e., parasite induced mortality) [8,9,27], parasites can also impact host fitness by altering host reproductive output [29–32]. Parasites can also indirectly influence host reproduction by reducing host body condition and available resources. This results in reduction of offspring number and\or quality [31]. Moreover, parasites can directly target the host’s reproductive organs to cause dramatic reduction of its reproduction up to castration [29,32,33]. Therefore, parasite virulence can be estimated also through its effect on host reproduction, especially when it is the main impact on host fitness [31,34–36]. Indeed, studies estimating the effects of coinfection on the reproductive success of the host demonstrated that, similarly to host survival, it can increase [20,37,38], reduce [20,39,40] or have no effect on it [41,42]. Since coinfections are common [1] and so is the negative influence of parasites on host reproduction [31], the effects of coinfection on host reproduction are probably common as well. Understanding these effects is crucial for obtaining a better comprehension of the impact of coinfections on host fitness. Furthermore, as many parasites are not lethal, the interaction between coinfections and host reproduction may play an important role in the regulatory function parasites have on host population density [43–45].

Here, we studied the fitness implications of coinfection by two parasite species of the water flea *Daphnia magna*, with emphasis on the reproductive component of fitness. *D. magna* is a common zooplankton inhabiting various aquatic ecosystems [46,47], where it plays an essential role in the local food-web [48,49] and the maintenance of water quality [50–52]. After spending approximately about two weeks in the juvenile phase, females mature and produce parthenogenic offspring with clutches of several to tens of neonates, that are released once every three days throughout their entire life [47]. In the field, *D. magna* is exposed to a variety of microparasites [46,53,54] and coinfections are commonly observed [55,56]. Moreover, the negative effect of coinfections on host reproduction [56] may play an important role in controlling *D. magna* population size [53,55]. Furthermore, reduced population density caused by parasites can be sufficient for generating a trophic cascade [57]. Therefore, the reproductive reduction induced by coinfections can potentially influence larger ecological processes in this system.

We used the microparasites *Metschnikowia bicuspidata* and *Hamiltosporidium tvaerminnensis* (hereafter, Met and Ht, respectively). Met is a common parasitic yeast that can infect various species of crustaceans including *D. magna* [47,58,59]. It resides in the host hemolymph and transmits horizontally after host death [47,60]. Ht (previously identified as *Octosporea bayeri* [61]) is a microsporidium that is currently known to infect only *D. magna* [47,62]. It is an intracellular parasite that infects the host’s adipose tissue and gonads, and can transmit both horizontally and vertically [47,63,64]. Both parasites can negatively affect host survival and reproduction [15,25,58,64]. While the effect of Met on the survival of *D. magna* is much stronger than that of Ht, the latter, especially in its vertical form, still dramatically reduces host reproduction [15,65,66]. Hence, the parasites differ in their mode of virulence, with Met mainly affecting host survival (about three weeks to host death [17,67]) and Ht mainly influences host reproduction. As these parasites naturally coinfect *D. magna* [55], they, along with the host, form a relevant triad with which to study the cumulative impact of coinfection on host reproduction.

We exposed *D. magna* to Met, Ht or both parasites in every possible order of arrival, including vertical transmission, and recorded how the interactions between the parasites affect the fitness of each organism. Our main objective was detecting how coinfection by parasites with different transmission and virulence strategies affects host reproduction. Our secondary objective was to describe the outcome of the interaction among the parasites. In previous studies, when parasites differed in their impact on host survival, coinfections were dominated by the more virulent parasite [25,68]. However, when coinfecting parasites had similar levels of virulence, neither parasite dominated and there was little to no additive effect on host survival [15,17]. Although host reproductive output in these studies exhibited a dominance effect [17], additive effect [15] and indifference [68], differentiating if host reproduction was directly or indirectly (via host mortality) affected was beyond their scope. In addition, a large variation in virulence was observed between Ht transmission modes [15]. Less virulent horizontal transmission was generally inhibited by the more virulent competitor, but both parasites suffered fitness loss when their virulence level became similar during vertical transmission. Hence, we predict that the balance of virulence among the parasites, influenced by Ht transmission mode will orchestrate coinfection’s outcomes. Therefore, host fitness components will be dominated by the mode of parasite virulence (i.e., survival for Met and reproduction for Ht). Alternatively, fitness effects of Ht will be impeded by drastic decreases in host survival caused by Met. For the parasites, we predict that the less virulent horizontal Ht will be outcompeted by Met. But when Ht is transmitted vertically, both parasites will suffer fitness loss, albeit asymmetrically, with some advantage to the more virulent Met.

## Results

Out of 500 *D. magna* individuals that started the experiment, five individuals were lost and excluded from the final dataset. In addition, two individuals from the groups that were vertically infected with Ht showed no signs of infection and were excluded from the dataset as well. We also excluded from the dataset 10 individuals that died before the age of 16 days (10 days post first parasite exposure), as we cannot identify visible signs of Met infection before that day. Therefore, we considered their death as background mortality. After these exclusions, the remaining dataset consisted of 483 individuals. From these individuals, 365 were infected, 71 were exposed but not infected, and 47 were control individuals. From the infected individuals, sixteen individuals were infected by Met but failed to develop mature spores and two individuals that were vertically infected by Ht and exposed to Met did not develop an infection with it. Hence, we excluded these individuals (n= 89) from the analyses of spore production, host survival and host reproduction along with the exposed but not infected individuals. More information about exposed but not infected individuals is available in the supplementary material. A summary of the final sample sizes and the distribution of infected individuals in the different treatments is available at Table 1.

**Table 1.** Experimental design and final sample sizes. Each control group is presented with its classification by exposure type and time of exposure. Under exposure type, ’vertical’ hosts were infected by Ht via vertical transmission (host age at exposure=0). Under early/late exposure, control-hosts were exposed to macerated *D. magna*; Met-hosts were exposed to 90,000 spores of *M. bicuspidata*; and Ht-hosts were exposed to 300,000 spores of *H. tvaerminnensis*. Under final sample size, numbers in round parenthesis represent all hosts infected by a parasite, including coinfection. Numbers in square parenthesis represent all coinfected individuals on the left side and animals that were infected by Met, but it failed to produce mature spores.

| Treatment | Exposure type | Early exposure<br>(Host age = 6 days) | Late exposure<br>(Host age = 11 days) | Final sample size (Infected by Met \| Ht) [Coinfected \| Met failed] |
| --- | --- | --- | --- | --- |
| Control | None | Control | Control | 47 (0\|0) [0\|0] |
| Met early | Single | Met | Control | 48 (31\|0) [0\|3] |
| Ht early | Single | Ht | Control | 49 (0\|24) [0\|0] |
| Met late | Single | Control | Met | 47 (47\|0) [0\|0] |
| Ht late | Single | Control | Ht | 49 (0\|46) [0\|0] |
| Ht + Met | Simultaneous | Ht + Met | Control | 50 (24\|4) [0\|1] |
| Ht + Met late | Sequential | Ht | Met | 49 (48\|0) [0\|0] |
| Met + Ht late | Sequential | Met | Ht | 48 (35\|14) [4\|4] |
| Ht vertical | Single<br>(vertical) | Control | Control | 49 (0\|49) [0\|0] |
| Ht vertical + Met | Sequential<br>(vertical) | Met | Control | 47 (45\|47) [45\|8] |

### Parasite infection rate

The presence of Ht (horizontal or vertical transmission) did not affect the probability of infection by Met (Table 2, Figure 1). However, time of exposure reduced infection probability by 30.7% in early exposure (Table 2, Figure 1). Comparison among the treatment groups nested in the early exposure revealed that the probability of infection by Met in individuals that were vertically infected by Ht was significantly higher than in all other groups (log odds ratio > 2, p < 0.01 for all pairwise comparisons, see supplementary material). While the probability of infection by Met in the presence of Ht horizontal spores differed by the time of exposure to Ht (simultaneous or sequential, log odds ratio = 1.059, p = 0.021), the success of Met in both treatments did not differ from the group that was not exposed to Ht (log odds ratio < 0.7, p > 0.1 for two comparisons).

**Figure 1.**
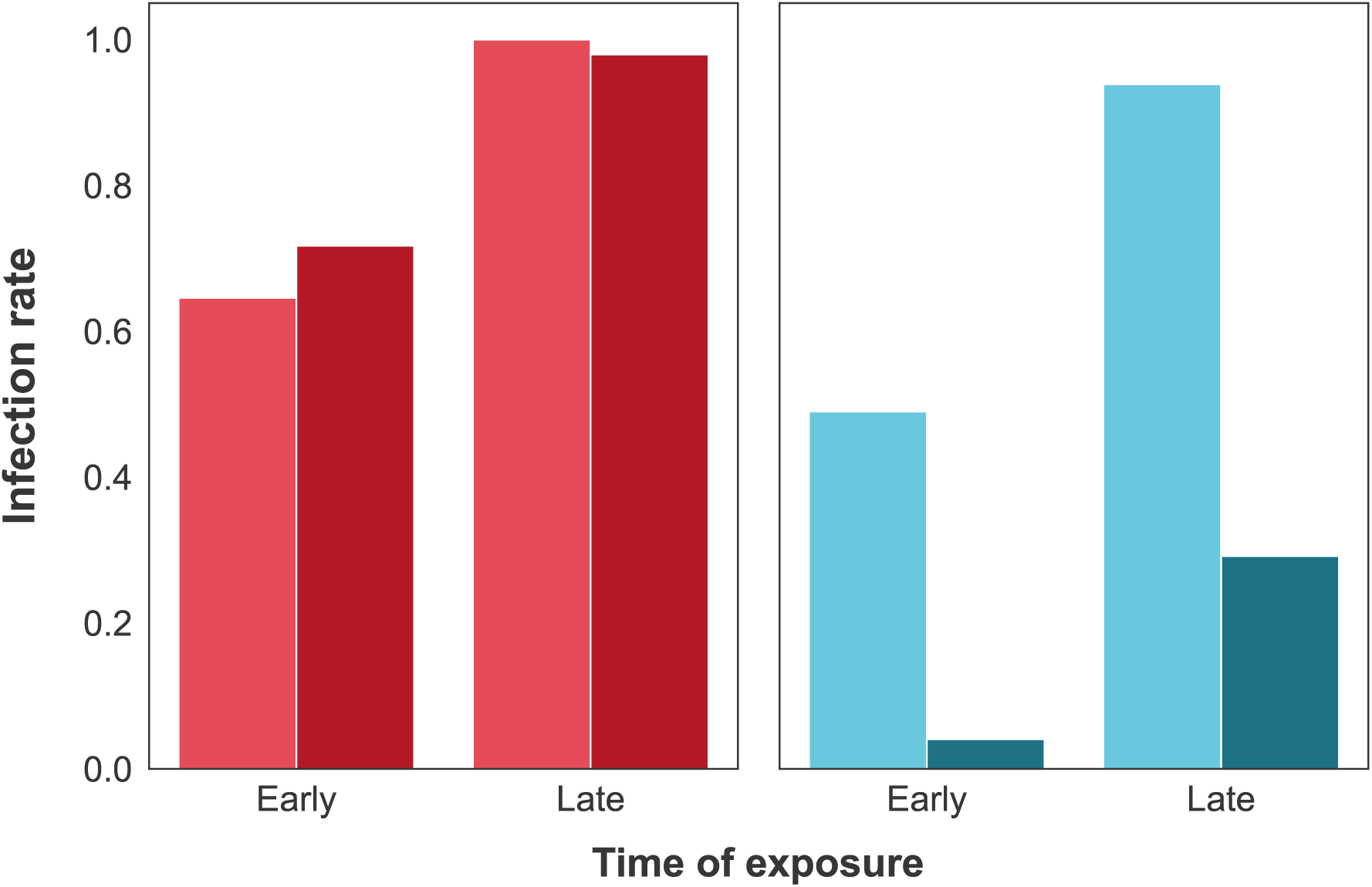
Infection rate of Met and Ht. Infection rates (represented by the proportion of infected individuals) for Met (red, left panel) and horizontal Ht (teal, right panel) are presented for early and late exposures. Lighter or darker shade of color represent parasite success when exposed to the host alone or with its competitor, respectively. Time of exposure but not the presence of competition significantly affected Met, while both factors significantly affected Ht. Additional details are available in Table 2.

**Table 2.**
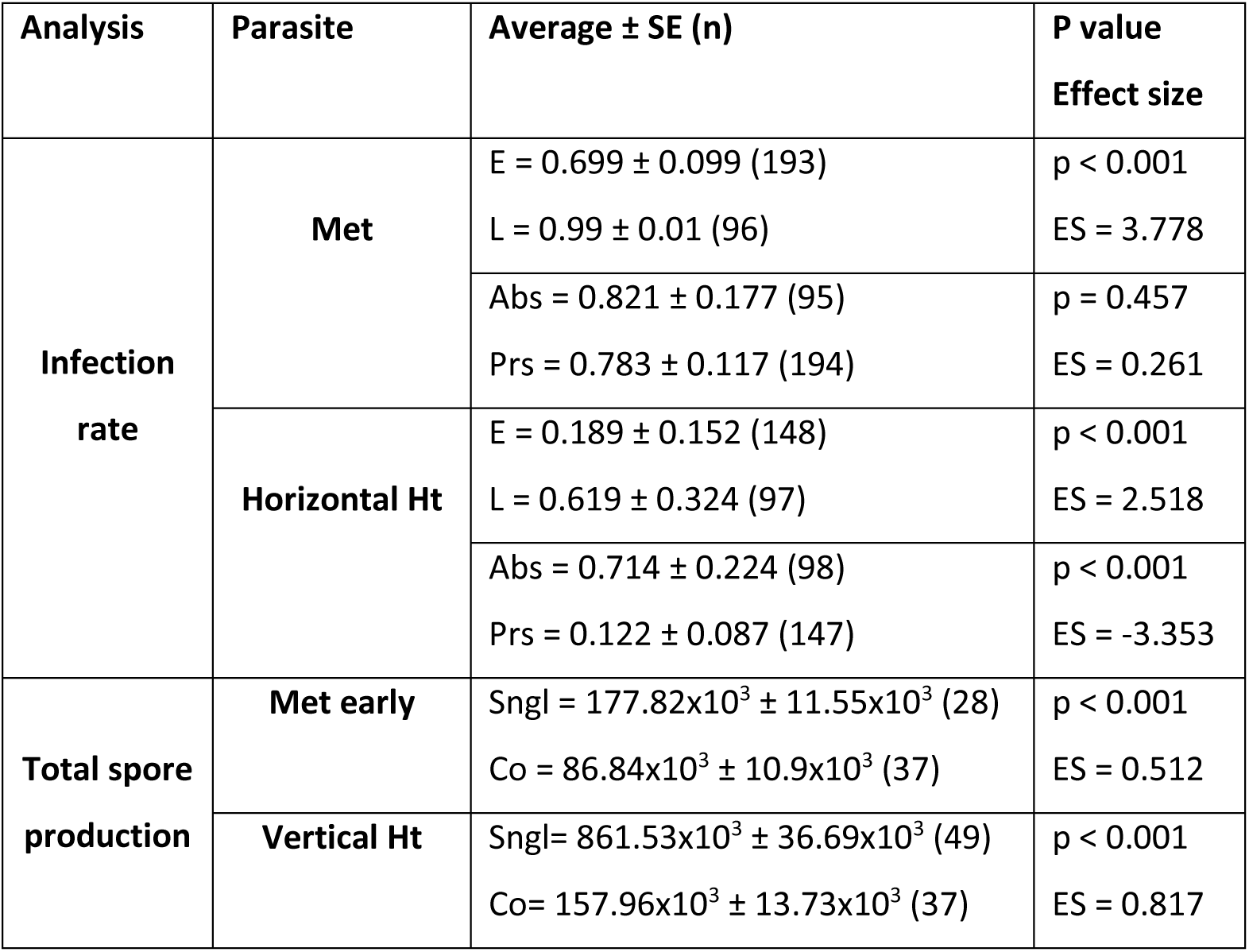
Effects of parasite competition on parasite success. Main effects of parasites on each other as indicated by the regression models. Averages and standard errors of infection probabilities and total spore production are presented for their respective analysis. Standard errors of infection probabilities were calculated from the variation among nested treatment groups, since it represents properly the variation in the observed data. Effect sizes of infection rate are estimated by log odds-ratios, while effect sizes of spore production are estimated by normalized differences of coinfection from single infection, i.e., (x_sngl_ – x_co_)/x_sngl_. Abbreviations: E=early exposure, L=late exposure, Sngl=single infection, Co=coinfection, Prs\Abs=competitor’s presence\absence.

Unlike Met, the horizontal Infection rate of Ht was significantly reduced in the presence of its competitor (Table 2, Figure 1). In addition, late exposure yielded a significantly higher infection rate than early exposure (Table 2, Figure 1). The additional multiple comparisons of treatments nested within factors’ categories revealed no deviation from these results: Met exposed groups always had lower infection rates regardless of time and late exposure always had higher infection rates compared with their peers regardless of Met presence (see supplementary materials).

### Parasite spore production

The effect of coinfection with vertically infected Ht resulted in Met losing approximately half of its progeny compared to single infection at the same exposure time (Table 2, Figure 2). Horizontal exposure to Ht had no effect on Met spore production nor did host age at infection (Table S3). Coinfection with Met resulted in even greater loss of spores for vertical Ht that lost about 80% of their progeny (Table 2, Figure 2). Once again, exposure that did not result in coinfection had no effect on spore production (Table S3). However, different host age at exposure resulted in major changes in spore production. The spore yield in vertical transmission (i.e., exposure at birth or pre-birth) was more than three times higher than in late horizontal exposure (host age of 11 days). The differences in both categories from early horizontal exposure (host age of 6 days) were by orders of magnitude (Table S3).

**Figure 2.**
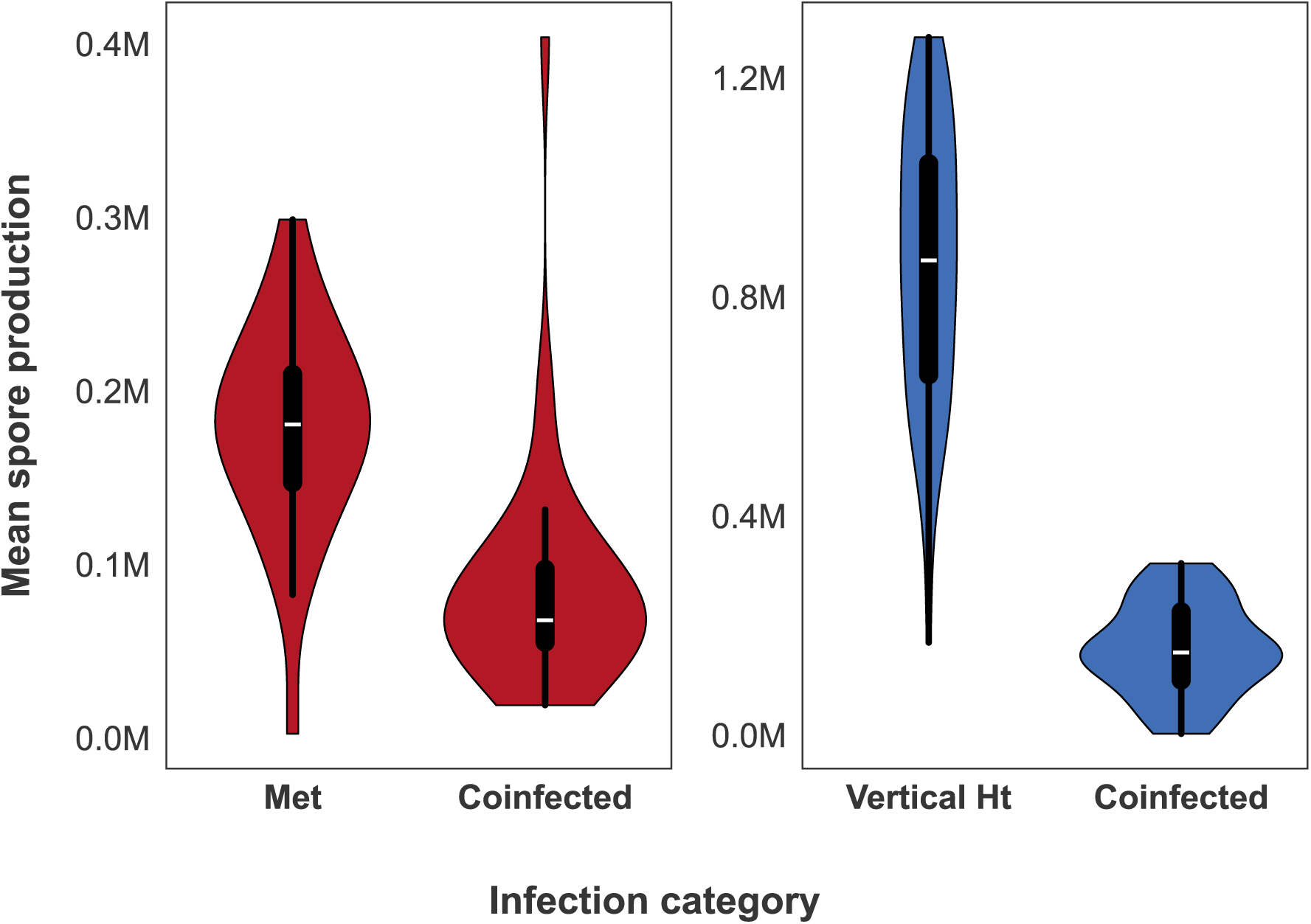
Parasites spore production under coinfection. Violin plots of total spore production (in millions) by Met (red. left panel) and vertical Ht (blue, right panel), when infecting the host alone or together with their competitor. The violin contours are proportional to the number of observations, while the data’s boxplots are presented inside. Spore production of the two parasites was significantly reduced under coinfection. Additional details are available in Table 2.

### Host survival

The structure of the final Kaplan-Meir model included the control, Met and coinfected categories, while the Ht category was split into vertical Ht, and early and late horizontal Ht (Figure 3, see supplementary material). All categories except for early horizontal Ht had a significant negative effect on host survival (Tables 3 and 4). Met-infected and coinfected individuals suffered more than tripled mortality rate than controls (Tables 3 and 4). These groups were similar, as reflected in the multiple comparisons and their non-significant log rank test in the final model (p=0.325). Among the Ht-infected groups, vertically-infected individuals induced medium reduction in host survival with almost double the mortality rate of controls. Late horizontal infections caused a small reduction in host survival together with a small increase in host mortality rate, but early infections had no negative effect and even survived slightly better than controls (Tables 3 and 4). All Ht-infected categories were significantly different from each other and from the Met and coinfected categories as well (p<0.001 for all log rank comparisons).

**Figure 3.**
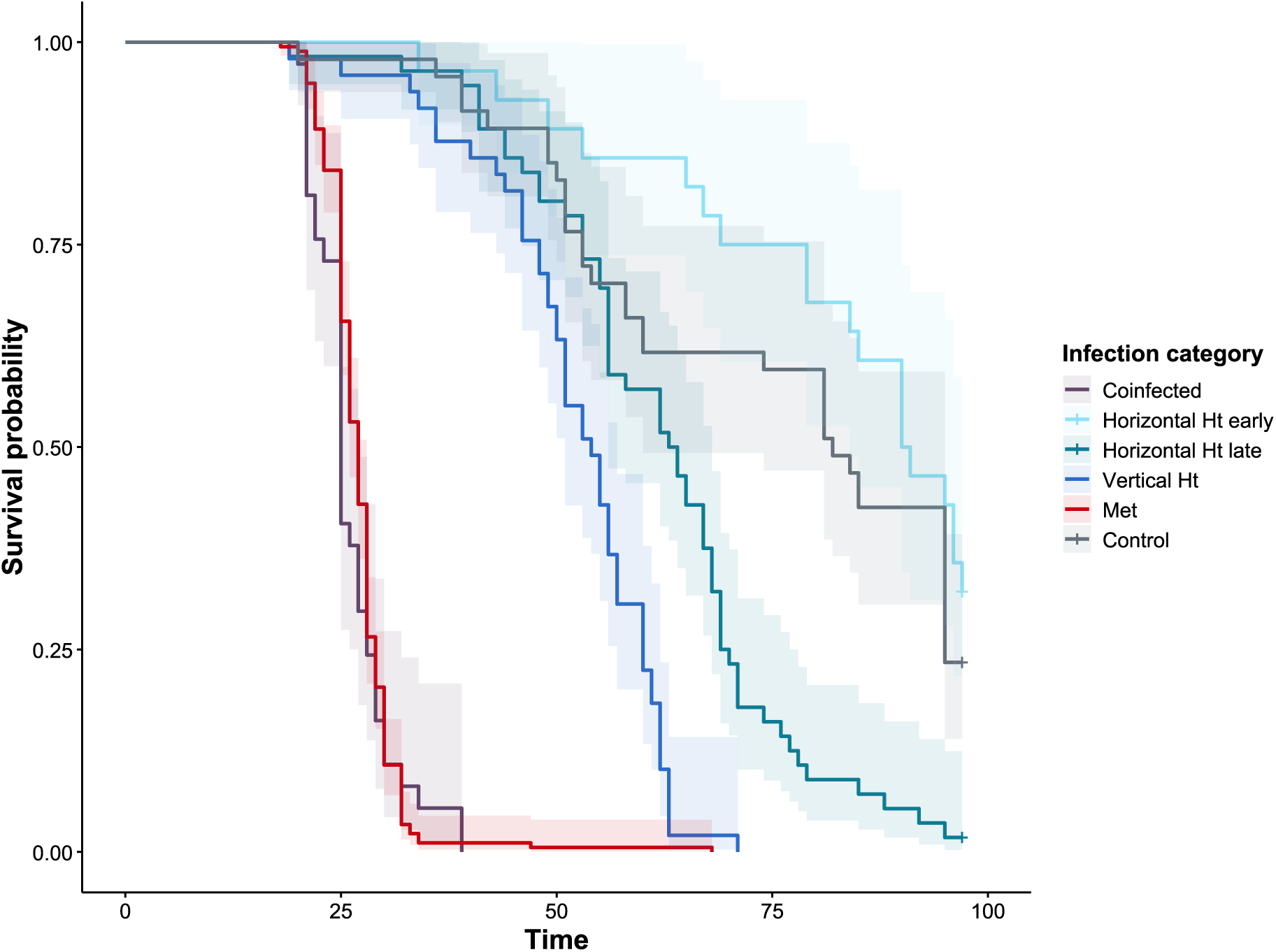
Host survival under different infection categories. The survival of hosts over time outlined by Kaplan-Meier analysis are presented for each infection category (Infection by Met – red, horizontal Ht – cyan and teal for early and late exposure, respectively, vertical Ht – blue, coinfected – purple, and control – grey). All infection categories except for early horizontal Ht were significantly different from the control category. Additional details are available in Tables 3 and 4.

**Table 3.** Host fitness under parasitic infection. Summary of host fitness parameters under different infection categories. Where relevant, results are presented in an average ± SE (n) format. ’Average’ indicates that the measurement for a single individual is the average of multiple events. Abbreviations: AUC=area under the curve, E=early, L=late.

| Fitness component | Parameter | Control | Horizontal Ht | Vertical Ht | Met | Coinfected |
| --- | --- | --- | --- | --- | --- | --- |
| Host survival | Longevity | 75.085 ± 3.316 (47) | E = 83.037 ± 3.549 (28)<br>L = 61.946 ± 2.041 (56) | 51.898 ± 1.5 (49) | 27.124 ± 0.346 (177) | 26.135 ± 0.749 (37) |
|  | AUC | 75.085 | E = 83.035<br>L = 61.946 | 51.898 | 27.124 | 26.135 |
|  | Mortality rate | -0.016 | E = -0.019<br>L = -0.021 | -0.033 | -0.053 | -0.057 |
|  | Weighted mortality rate | -0.008 | E = -0.007<br>L = -0.01 | -0.014 | -0.015 | -0.026 |
| Host reproduction | Age of first event | 16.34 ± 0.351 (47) | 15.286 ± 0.204 (84) | 16.108 ± 0.657 (46) | 14.585 ± 0.144 (176) | 14.29 ± 0.636 (31) |
| | Average interval between events | 4.105 ± 0.078 (46) | 3.865 ± 0.047 (83) | 4.856 ± 0.183 (45) | 3.973 ± 0.129 (172) | 6.818 ± 0.676 (22) |
| | Average clutch size | 13.196 ± 0.315 (47) | 11.782 ± 0.247 (84) | 8.215 ± 0.399 (46) | 6.315 ± 0.156 (176) | 3.28 ± 0.365 (31) |
| | Clutch size adjusted for time * | 10.06 ± 1.412 (47) | 8.87 ± 1.237 (84) | 5.71 ± 0.807 (46) | 7.93 ± 1.111 (176) | 4.82 ± 0.774 (31) |
| | Total offspring produced within first 39 days of life | 193.638 ± 10.355 (47) | 168.81 ± 6.965 (84) | 72.152 ± 5.092 (46) | 23.358 ± 1.193 (176) | 5.677 ± 0.576 (31) |

\* Averages and SEs are the adjusted marginal means calculated from the model to account for the effect of time.

**Table 4.** Effects of parasites on host fitness. Effect sizes of each infection category in relation to the control group and p-values of their paired comparison. All survival parameters were derived from the same analysis, hence their p-value is presented once. Unless stated otherwise, all effect sizes were estimated by the normalized difference from control. Effect sizes of age on first events were estimated using the raw difference from control and those of mortality rates were estimated by the ratio infected by control. Abbreviations: AUC=area under the curve, E=early, L=late.

| **Fitness component** | **Parameter** | **Horizontal Ht** | **Vertical Ht** | **Met** | **Coinfected** |
| --- | --- | --- | --- | --- | --- |
| **Host survival** | **AUC** | E:<br>p = 0.289<br>ES = -0.106<br>L:<br>p < 0.001<br>ES = 0.175 | p < 0.001<br>ES = 0.309 | p < 0.001<br>ES = 0.639 | p < 0.001<br>ES = 0.652 |
|  | **Mortality rate** | E = 1.14<br>L = 1.274 | 1.986 | 3.229 | 3.463 |
|  | **Weighted mortality rate** | E = 0.886<br>L = 1.282 | 1.784 | 1.862 | 3.247 |
| **Host reproduction** | **Age of first event** | p = 0.042<br>ES = 1.055 | p = 0.948<br>ES = 0.231 | p < 0.001<br>ES = 1.755 | p = 0.002<br>ES = 2.05 |
|  | **Average interval between events** | p = 0.399<br>ES = 0.942 | p = 0.029<br>ES = 1.183 | p = 0.551<br>ES = 0.968 | p < 0.001<br>ES = 1.661 |
|  | **Average clutch size** | p = 0.001<br>ES = 0.107 | p < 0.001<br>ES = 0.378 | p < 0.001<br>ES = 0.519 | p < 0.001<br>ES = 0.751 |
|  | **Clutch size adjusted for time \*** | p < 0.001<br>ES = 0.118 | p < 0.001<br>ES = 0.432 | p < 0.001<br>ES = 0.212 | p < 0.001<br>ES = 0.521 |
|  | **Total offspring produced**<br>**within first 39 days of life** | p = 0.171<br>ES = 0.128 | p < 0.001<br>ES =<br>0.627 | p < 0.001<br>ES =<br>0.879 | p < 0.001<br>ES = 0.971 |

\* Calculated from the adjusted marginal means and estimated from the model to account for the effect of time.

### Host reproduction

All infected categories experienced a significant reduction in the average clutch size compared to the control group (Table 4). The largest reduction was in the coinfected group, with a 75% reduction (Tables 3 and 4). Additional strong negative impacts were of Met and vertical Ht infections that reduced the average clutch size by approximately 52% and 38%, respectively, whereas the effect of horizontal Ht was much smaller (about 11%; Tables 3 and 4). However, the impact of parasites on host reproduction changed when the effects of host age on clutch size was taken into account (Tables 3 and 4 as well as supplementary material). Reanalyzing the effect of parasites on *Daphnia* clutch size while controlling for the effect of time demonstrated that vertically transmitted Ht (alone or in coinfection) caused the greatest reduction. This form of Ht reduced up to 52% of host clutch size throughout its life (Tables 3 and 4, Figure 4). The effect of Met, on the other hand, was much smaller, while the effect of horizontally transmitted Ht remained the same (Table 4). The actual difference in estimated clutch size of both Met and horizontal Ht from the control was between 1-2 offspring (Table 3). Furthermore, analyzing the AUC differences along the dynamic of host reproduction through time stressed the differences between the effects of vertical Ht and the other categories. Animals infected vertically with Ht (including coinfected animals) had a large, consistent and significant negative effect on their clutch size throughout their lifespan (Figure 4). On the contrary, the effect of horizontal Ht and Met was limited in time, and usually small, as both parasites limited their host clutch size at the peak of infection (trend at the first and significantly at the second, Figure 4). The difference between these two categories was that Met caused another significant reduction between weeks 4 and 5 (Figure 4). It is important to note that the sample size of Met from week 7 onwards was very small (Figure 4). Though we would usually omit samples of this size, since relatively long lifespan under Met infection is rare, we find these surviving individuals as an illustration of the reproductive potential under these circumstances, albeit interpret these findings with caution.

**Figure 4.**
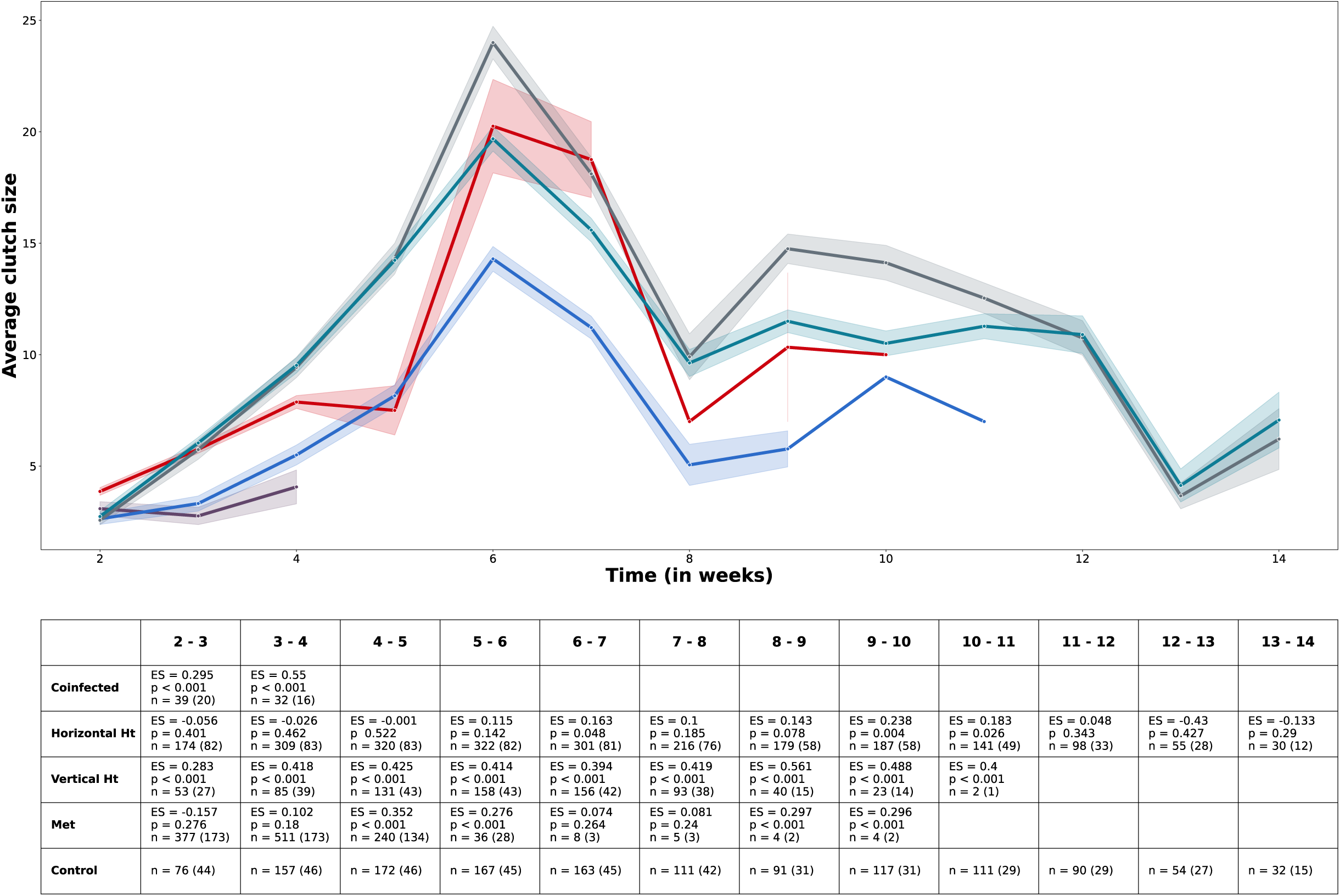
Host reproduction under different infection categories over time. The average number of offspring in a single clutch during the lifetime of hosts is presented (top panel) for each infection category (infected by Met – red, horizontal Ht – teal, vertical Ht – dark blue, coinfected – purple, and control – grey). The table in the bottom panel presents the effect sizes and p values from the AUC analyses, respectively. Each column refers to an AUC of two consecutive weeks and the effect sizes are the normalized differences of each treatment group from the control. In addition, the sample sizes of each timeframe in the analysis are by both the total number of clutches released and the maximal number of reproducing individuals (in brackets). The maximal number of reproducing individuals can be smaller than the number of individual alive in the respective time frame, if some individuals did not reproduce, e.g., coinfected individuals that survived beyond the fourth week were completely castrated and are not represented in neither the figure nor the table. In general, hosts produced more offspring pre-clutch up to their sixth week of life and gradually decreased offspring production towards the end of the experiment. While this trend was not different between groups, parasites reduced the number of offspring produced to a varying degree, but only the negative effect of vertical Ht (including coinfected hosts) was consistent throughout the host’s life.

Animals infected horizontally tended to exhibit early onset of reproduction. In particular, those infected by Met (including coinfection) started to reproduce two days before control animals, while those infected by Ht started to reproduce one day earlier. Vertical infection with Ht had no effect on the onset of reproduction (Tables 3 and 4). In addition, coinfected individuals had 1.66 times longer interval between clutches, whereas other groups had no or much smaller intervals (Table 4).

Finally, comparing total host reproductive output during the first 39 days illustrated how all effects above come to conclusion. More precisely, in comparison to control, coinfected individuals lost almost all their reproductive output (approximately 97%, Tables 3 and 4). Though having a small direct effect on host reproduction, Met-infected individuals suffered as well from a dramatic reduction of about 88% due to their short lifespan (Tables 3 and 4). Individuals infected vertically with Ht lost 63%, evidence of the strong impact of the parasite on host clutch size. Lastly, the total output of *Daphnia* infected horizontally by Ht was not significantly different from that of controls (Table 4).

## Discussion

Using *D. magna* as the host, we studied how coinfection by two parasites that differ in their transmission mode and virulence affects host reproduction and survival. We found that coinfected individuals suffered from greater reduction in reproduction but not in survival. We demonstrated that the effect of vertical Ht on host clutch size was greater than the effect of Met and that it dictates this trait during coinfection just like Met dictates host survival. While this result meets our prediction, the excess reproductive loss of the host cannot be explained solely by adding the effects of the two parasites. Synergetic effects of coinfection reduced the number of clutches released by the host and led to severe reduction in the lifelong reproduction of coinfected hosts. These effects demonstrate how increased virulence experienced by the host can occur via its reproduction, beyond the additive effects of coinfection.

Host experience under coinfection is often unique and cannot necessarily be deduced from the additive expectation of single infections by the coinfecting parasites [10,18,69]. For example, mice suffered from high mortality when coinfected by two species of rodent malaria that would not be expected from the survival rates observed in single infection by the same species [69]. In our study, host reproductive output under coinfection was significantly lower compared to single infection within the timeframe of coinfected host longevity. This effect could not be expected from the joint direct effect of Ht on host clutch size and the indirect effect of Met-induced mortality. Two unique effects that were not observed in single infection contributed to the excess reproductive loss: the interval between reproductive events was nearly doubled and the host became castrated if it survived beyond the fourth week mark of the experiment. Hence, coinfected hosts not only produced smaller clutches due to Ht infection, but also produced fewer of them. While many studies justifiably focused on the implications of coinfection on host survival, our results as well as others [15,20,39] suggest that studying multiple components of fitness is necessary for comprehensively grasping host experience under coinfection. Such an approach is highly relevant for iteroparous r-selected hosts, as their reproductive success does not completely depend on surviving until a specific day of offspring release. *Daphnia* reproduce frequently after reaching sexual maturity and can produce numerous offspring during their lifetime under lab conditions [47]. However, in the field, individuals may not be expected to fulfill such reproductive potential, as they are predated on by numerous organisms [57,70,71] and exposed to virulent parasites [53,72,73]. Thus, every change in immediate reproductive success may have a larger than expected influence on their population density and genetic structure. Considering the variation in the capacity of parasites to induce host mortality [47,58], changes in host reproduction mediated by coinfection are equally important as changes in host survival. In addition, the potential consequences for hosts that manage to survive aggressive infections [74–76] can reveal reproductive loss as another negative consequence for populations exposed to virulent parasites. Therefore, additional emphasis on the implications of coinfection on host reproduction should generate a more holistic perspective on the ramifications of this process for wild populations.

The outcome of coinfection is the product of interactions among the parasites and their host [8,14,38,77]. Depicting the network of interactions based on the outcome of coinfection is complicated, since multiple processes can operate simultaneously, at different levels and in various directions to shape the outcome [14]. Yet despite their complexity, studies were able to outline them [78], find their relative commonness [79] and frame them within known ecological rules [14]. For instance, bottom-up processes, such as exploitative and interference competition among parasites, are expected to reduce parasite fitness [8,14,77]. Additionally, the relative effect can be also shaped by the level of asymmetry of these interactions. For example, Massey et al. [8] demonstrated how maximal asymmetry between coinfecting bacteria species (one can inhibit the other but not vice versa) resulted in exclusion of the subordinate while host fitness followed the winer. However, when the coinfecting species had similar influence on each other (smaller asymmetry), it resulted in reduced fitness for both species while the host experienced reduced virulence. In our study, when both parasites were introduced horizontally, Ht was generally excluded and host fitness was determined by the winner (usually Met). Current molecular knowledge links the virulence of Met to within-host growth and exploitation [80,81]. This suggests that our observed exclusion is probably the result of exploitative mechanisms. Fast expansion by Met may have left little time and/or resources for the seemingly slower Ht. In addition, attaching to host cells is crucial for microsporidian establishment within the host [82]. The fast-expanding hyphae of Met may have blocked or reduced Ht’s adhesion and indirectly inhibited its establishment. The mutual reduction in parasite spore production under coinfection further supports this direction. Several studies have found that the availability and quality of resources is positively linked with the reproductive success of *Daphnia* microparasites [83–86]. Moreover, *Daphnia* parasites can change the resource usage of their host [66,87] and even reduce their nutritional values for predators [88]. Hence, it is likely that the reproductive loss of our parasites was mediated through resource exploitation.

Resources competition among our parasites can also explain the excess reproductive loss of the host. *Daphnia* eggs are relatively rich in nitrogen [87] and changes in the use of nitrogen were found to be correlated with fecundity loss under parasitism [66]. Furthermore, the spores of one *Daphnia* parasite, *Pasteuria ramosa,* were found to be rich in nitrogen, though less than host eggs [87] and Met produced more spores in the presence of nitrate [89]. While parasites can differ in their use of key nutrients [66], if nitrogen is an important component for spore production, our result could be the product of excess pressure by two reproducing parasites. Additionally, Met’s impact on host reproduction is focused towards the end of the host’s life (Figure 4), when the burden of spore production is at its peak. This period parallels the period during which coinfected hosts experienced castration. This may suggest that it is not just that the damage to the host intensified over time, but also that resource competition could have escalated with time. Still, our study cannot rule out contribution from additional types of interactions (e.g., host immune response) to our results. Nonetheless, the mechanistic approach has been suggested to be advantageous for the identification of interactions among organisms [90] and assist in the classification of parasite function within the host [91], which can improve the predictability of coinfections [92]. Improving our abilities to directly link between active mechanisms and coinfection outcomes is expected to contribute beyond the realm of ecology and evolution.

In addition to the importance of parasite-parasite interactions, various factors related to the interacting organisms [7,9] and the environment [10,11] can further influence parasite fitness. Here, parasite success was mainly affected by a single factor: the transmission mode of Ht. When transmitted horizontally, Ht was in general competitively excluded by Met (Figure 1), but became a better competitor when transmitted vertically, which resulted in significant loss of spores for both parasites (Figure 2). One may argue that vertical transmission is simply another type of priority effect affecting parasite success in coinfections [23,93,94]. However, none of the results we presented here are explained better by the timing of infection rather than Ht’s transmission mode. The exclusion of horizontal Ht was not affected by the order of arrival nor was its reproductive output that matched that of single exposure controls (Supplementary materials). Likewise, the increased infection rate of Met is likely due to a stochastic rather than ecological process (Supplementary material). More importantly, the infection profiles of the two transmission modes were extremely different. Vertical transmission was more virulent when investment in horizontal transmission increased (see additional discussion in the supplementary material). Similarly, different developmental stages of the trematode *Echinostoma caproni* demonstrated different infection profiles in both virulence and reproductive strategy [42]. These in return greatly influenced the coinfection outcome with *Schistosoma mansoni* within the same host species [20,42,95]. Therefore, it is reasonable to interpret the different modes of Ht as different developmental stages that interact differently with Met according to the infection profile. The consistency of our findings with those of Ben Ami et al. [15] suggests that the effects of Ht’s transmission mode on its success in coinfections is a true characteristic of this parasite.

We acknowledge that our single parasite isolate-single host maternal line design could have missed variations in the final outcome that can result from host-parasite or parasite-parasite GxG interactions [9,96][7]. Nonetheless, genetic variation seems to produce variation around a general trend that is consistent between the infecting species [9,96]. Furthermore, our results were generally consistent with those of Ben Ami et al. [15] that used different genetic lines of *D. magna* and Ht and a different coinfecting parasite species (the castrating semelparous bacterium *P. ramosa*).

To conclude, coinfection by *M. bicuspidata* and *H. tvaerminnensis* increased the virulence experienced by their *D. magna* host. However, the increased virulence was not due to increased parasite-induced host mortality, but through decreased host offspring production. As the lifetime reproduction of coinfected individuals was, on average, less than a single clutch in comparison to uninfected hosts, our results advocate to dedicate more attention to the effects of coinfections on host reproduction. Moreover, such a strong effect may speed-up evolutionary ’Red Queen’ dynamics [97] in natural populations by further reducing the contribution of coinfected individuals to the gene pool of the next generation. Since coinfections can be random [55,98], they can even the evolutionary playground if reproductive reduction acts regardless of the identity of the coinfecting parasites. Under this scenario with frequent coinfections, many individuals may suffer, to varying degrees, reproductive losses that can result in more complex and slower genetic dynamics. Therefore, the interplay between coinfections and evolutionary dynamics should be an interesting endeavor for both theoretical and empirical studies.

## Materials and methods

### Experimental design

We randomly allocated 50 females of *D. magna* from a single maternal line to one of ten treatments (Table 1). While all individuals originated from the same host culture (see below), the individuals of the two treatments involving vertically transmitted Ht originated from a culture of the same maternal line vertically infected with Ht at ∼100% prevalence. All treatments were exposed to parasite spores suspensions or to a control solution when they were six and eleven days old. The type and order of exposures of each treatment is detailed in Table 1. The experiment was terminated 91 days after the first exposure.

### Organism cultures

Two populations of *D. magna* females of the same age were kept in groups of 10 and placed in 250 ml glass jars filled with ventilated tap water. The individuals were transferred to new jars twice a week to separate adults from their offspring. While the two populations were of the same maternal line one population was kept uninfected, while the second was comprised of individuals that were vertically infected with a single isolate of Ht to enable the collection of vertically infected individuals. In addition, we cultivated dense populations of our *Daphnia* infected with the same isolate of Ht or with a single isolate of Met. These infected populations were kept in ventilated tap water with additional water added to the jar, when necessary, to prevent loss of parasite spores. All populations were fed daily with sufficient amounts of the green algae *Scenedesmus sp.* and were kept at a temperature of 20° with 12:12 light:dark cycle. Populations infected with Met were occasionally supplemented with ten uninfected PS-2 individuals to prevent their extinction.

### Infection assays

PS-2 offspring from both uninfected and Ht infected populations born within 24 hours were collected and kept in low density groups for 3 days, and fed daily with a sufficient amount of *Scenedesmus sp.* as previously described. On day 4 all offspring were sexed under a dissecting microscope (Leica M205C) and females (400 uninfected and 100 vertically infected with Ht) were allocated individually to glass jars filled with 20 ml of ventilated tap water. Two days later, on day 6, each individual was exposed to 90,000 spores of Met, 300,000 spores of Ht, spores of both parasite species, or a control suspension made from uninfected PS-2 individuals that were macerated in ventilated tap water according to their treatment group (early exposure in Table 1). Feeding was skipped on the day of exposure to increase the *Daphnia* filtration rate. Forty-eight hours post exposure, all individuals were placed in a clean jar with 20 ml of ventilated tap water. Five days post exposure, on day 11, the individuals were exposed to spores or control suspensions according to their treatment group (late exposure in Table 1). While the suspensions were made fresh, the number of spores individuals were exposed to was the same as in the early exposure. Similarly, feeding was skipped on the day of exposure and after 48 hours all individuals were placed in clean jars, this time filled with 100 ml of ventilated tap water. Once a week, all individuals were transferred to clean jars with 100 ml of ventilated tap water. Except for exposure days, individuals were fed every day except on Saturday following an age-dependent feeding protocol.

### Host reproduction and survival

From the first exposure until the end of the experiment, individuals were monitored daily (except on Saturday) for reproduction and survival. Offspring were removed from the jar and their number was recorded. Dead individuals were placed in an Eppendorf tube filled with 20% glycerol solution, stored in −20°C and their date of death was recorded. On the last day of the experiment, all surviving individuals were preserved in the same way. For survival analysis (see below), we considered this date as the death date of the surviving individuals, but flagged them as censored.

### Parasite infection rate and spore production

Every host was defrosted and macerated to a homogenous solution. An aliquot of the solution was placed on a hemocytometer (Neubauer new) and spores were counted under a phase-contrast microscope (Leica DM500). When a sample was too dense, it was diluted with 20% glycerol solution and the new volume of the tube was recorded. The counts were converted to concentrations [104] and total spore production was estimated by finding the product of the solution’s concertation and volume. To maintain consistency, a single observer counted the spores of each parasite. For Met, only mature spores were counted (following illustrations no. 16-19 in [105], while the presence of spores in other stages of their development was sufficient to count the individuals as infected without reproductive output. For Ht, all shapes of spores [64] were counted, as they were all considered infective [106].

### Data analysis

Since the analyses were mainly focused on infection processes and their consequences, except for the infection rate analyses, all exposed animals that did not develop infection were excluded from the analyses along with individuals that were infected by Met but failed to produce any mature spores. Additional information on final sample sizes, including specific analyses, is available in Tables 1, 2 and 3.

Parasite fitness was estimated based on their infection rate and spore production. To test for the effect parasites had on each other’s infection rate, we performed logistic regression for each parasite species separately. We used the infection rate of the target parasite as the dependent variable and exposure to the competing parasite as an explanatory variable. Since the date of exposure could have an effect on infection rate due to host age effects [107,108] or stochasticity, we added a categorical variable, ’exposure time’, representing the exposure dates as their relative position in the experiment’s timeline (i.e., early and late for first and second exposure dates, respectively). Since some treatment groups resulted in a variance of zero, we did not add an interaction term to the model. If any of the explanatory variables yielded significant result in the model, we performed multiple pairwise comparisons between groups nested within each category of the variable. We used Fisher’s exact test for the pairwise comparisons and controlled for false discovery rate at the 0.05 level when multiple comparisons were made. For Ht, the analysis included only the horizontal transmission treatments, since vertical transmission treatments were comprised only from infected individuals. For similar reasons, we did not analyze the coinfection rate. In the vertical transmission treatment, coinfection rate is equal to Met infection rate and coinfection in the horizontal treatments occurred only in four individuals (Table 1).

Similarly, we used linear regressions to identify the factors affecting parasites spore production while focusing on the effect of coinfections. We examined the effect of coinfection on the vertically infected Ht group that was exposed to Met with the relevant comparison group for each parasite (see Met early and Ht vertical in Table 1). This is because coinfections were common only in *Daphnia* that were infected vertically with Ht. Additionally, rare cases of coinfection with horizontally infected Ht (n=4) occurred only when Met failed to develop mature spores and therefore, they were excluded from the analysis. The model included the total amount of spores as the dependent variable and coinfection (binary) as the explanatory variable. Since many individuals infected by Met were exposed to, but not infected by, Ht, we used another linear regression to test if this condition influenced spore production. Like in the previous model, the total amount of spores was the dependent variable and exposure to Ht was the explanatory variable. We added ’exposure time’ to the model (in the same fashion mentioned above) to account for potential differences due to different exposure dates. For Ht infected individuals, only fourteen individuals were exposed to, bot not infected by Met. It resulted in unbalanced data structure that also violated the assumption of normality; hence, we split the analysis to two. First, we used a linear model to test for the effect of exposure date (including vertical transmission) on Ht’s spore production excluding the Met exposed individuals. Second, we used the Mann-Whitney U test to test for the effect of Met exposure on Ht’s spore production.

The analyses of host fitness were focused first on the effects of the different parasite species (when infecting alone or in coinfection) on host survival and reproduction. Therefore, each analysis started with a general model pooling all infected individuals, assigning them to one of three categories (Ht, Met or coinfected) and comparing them to the control group. Then, we tested within groups to determine if factors that affected parasite success created significant variation in the groups that affected their hosts. We used Kaplan-Meier survival analysis to test how different infection categories affected host longevity. A general model was constructed with individual longevity from birth as the survival index with an additional index marking animals that survived to the end of the experiment and used the infection categories as a stratifying factor. Then, we used the log-rank test for multiple paired comparisons between categories and controlled for a false discovery rate at the 0.05 level. In addition, we estimated the effect size of the different groups in two ways. First, we compared the difference in survival potential throughout the experiment by computing the area under the survival curve (AUC) and comparing the relative difference of these areas from the area of the control group. Second, we compared the mortality rates of each group and their relative deviation from the control group. We calculated both the average rate and the weighted average rate, since the first is more sensitive to the number of events and represents better the intensity of mortality, while the second is more sensitive to the length of events, represents differences that occur late on the timeline and creates a better predictor for future survival of individuals with extreme longevities. For more details on the calculation and rationale for choosing these parameters, please see the supplementary material.

We used a similar line of inquiry to estimate how our parasites shape the reproductive output of their host. Our starting point was to evaluate the effects of Met, Ht and coinfected categories on average clutch size. To estimate differences among categories and their nested groups, we used linear regression models (average clutch size of an individual and infection category as dependent and explanatory variables, respectively) with post-hoc multiple comparisons, controlling for false discovery rate at the 0.05 level. Since clutch size is a central characteristic of reproductive output, we assumed that the significant categories identified in this analysis would be the important categories to test for in subsequent models. Hence, we used the category structure from the average clutch size analyses (control, Met, horizontal Ht, vertical Ht and coinfected) for all the following models without further testing any nested groups. The next step of the reproductive output analysis was to isolate the direct effects of parasites on their host reproduction. If a parasite shortens host lifespan, then it is possible that any loss of progeny is a side effect of host mortality. This is especially true if clutch size depends on host age and changes over time. To this end, we tested for the effect of age on the clutch size of the control group, and determined if clutch size dynamics are different between infection categories. We then used generalized mixed model with an individual’s identity and age (in weeks) as random effects to test the effects of parasites on host clutch size when it is controlled for host age. Lastly, we extended the resolution of this model and analyzed the strength of parasites’ effects over time. We estimated the normalized AUC differences between the reproduction curves of the infected categories against the control in weekly intervals, and used a bootstrap-like simulation of virtual control populations to evaluate the significance of the observed parasites’ effect sizes. Additional details on these models are provided in the supplementary material.

We analyzed two additional aspects of reproduction. First, we asked if different infection categories affect the age a *Daphnia* released its first clutch. We used linear regression with age at the first clutch as the dependent variable and infection category as the explanatory variable. Second, we asked if there is an effect on the average interval between reproductive events. We used a generalized linear model (gamma distribution) with the average interval length as the dependent variable and infection category as the explanatory variable. To conclude this part, we analyzed the differences in sum of offspring from birth up to the maximal survival of the coinfected group (39 days; the first category to die out). While lifelong reproduction will produce larger differences, we believe that our timescale is a more relevant comparison. This is because the direct damage to reproduction takes place when the animal is alive, especially for *Daphnia* that is preyed by many organisms in their natural habitats [57,70,71] and thus less likely to achieve the longevity observed in the lab. Like in previous models, sum of offspring was used as the dependent variable and infection category as the explanatory variable. All the analyses were followed by post-hoc analyses of multiple pairwise comparisons controlled for false discovery rate at the 0.05 level. Finally, we tested if exposure to parasites that did not result in infection affected host fitness. Further details on these analyses and their results are available in the supplementary material.

All analyses were made in R version 4.2.2. We used the base package for the LM, GLM, Fisher exact test and the Mann-Whitney U test. For LMM and GLMM, we used the lme4 package [109] and for survival analyses we used the survival [110] and the surveminer [111] packages. Unless mentioned differently, we used the emmeans package [112] for all post-hoc pairwise comparisons. We produced the survival figures with the ggsurvfit [113] package and for the rest we used Python version 3.9, mainly with the seaborn [114] and matplotlib [115] modules.

## Supporting information

Supplemntary information and results

## Acknowledgments

A special thanks is dedicated to Gili Rotbard for her valuable assistance in the pre-liminary lab work. We also thank Yair Wexler for his valuable advices on the statistical analysis, Sofia Paraskevopoulou and Alex Dorfman for brain storming and Elizabeth M. Warburton for her useful comments on the first version of the manuscript.

## Authors’ Contributions

SH: conceptualization, methodology, investigation, methodology, project administration, data curation, formal analysis, visualization and writing- original draft; AS: investigation and data curation; FBA: funding acquisition, conceptualization, methodology, supervision and writing- review and editing.

All authors gave final approval for publication.

## Ethics

This study adheres to local guidelines. No ethical approval was required to conduct this study.

## Use of Artificial Intelligence (AI)

We declare that there was no use of AI in the creation of this manuscript.

## Data Availability

The datasets supporting this article as well as the R and Python script used for the analysis and visualization of this data will be on Figshare after formal publication of the manuscript. Any questions or requests can be sent to SH at.

