## Supplementary material for "Counterproductive: coinfection of a water flea by a fungus and a microsporidium reduces the reproductive outputs of all parties": Supplemntary information and results

##### PARASITE SUCCESS – ADDITIONAL INFORMATION

***Infection rate of Met in groups nested within exposure times***
**Table S1.**

***Infection rate of Ht in groups nested within exposure times or within presence\absence of competitor***
**Table S2.**

***Spore production – additional results***
**Table S3.**

***Differences between Ht transmission modes in single infection***

##### HOST SURVIVAL ANALYSIS – ADDITIONAL INFORMATION

***Calculations of effect sizes for Kaplan-Meir models***

***Results of the base model and follow-up comparisons***

##### HOST REPRODUCTION – ADDITIONAL INFORMATION

***The effect of parasites on host reproduction in the perspective of time***

***Results of the base model and final infection category's structure***

***The dynamics of clutch size and host age***

##### EFFECTS OF EXPOSURE TO PARASITES THAT DID NOT RESULT IN AN INFECTION
**Table S4.**

### Parasite success – additional information

#### *Infection rate of Met in groups nested within exposure times*

We compared the infection rates of treatment groups within each exposure event with Fisher's exact test. Early exposure was comprised of four groups, and we controlled for false discovery rate at the 0.05 level. The groups were single exposure, dual exposure with Ht spores, both simultaneously and sequentially (Ht second), and exposure to vertically-infected hosts (full details are provided in Table 1 in the main text). The comparisons revealed differences between groups with Ht exposure. The infection rate in vertically-infected host was like the rate during late exposure and significantly differed from all the other groups (Table S1). The simultaneous and sequential groups differed from each other (higher rate in the sequential group, Table S1), but did not differ from the single exposure group. In addition, the effect size of this comparison was at least half that of their comparisons to the vertically-infected group (Table S1). Late exposure was comprised of two groups, single exposure and sequential (Ht first, Table 1 in the main text). Both groups yielded high infection rates and did not differ from each other (Table S1). Interestingly, the increased success of Met to infect vertically infected individuals may suggest it benefits from Ht's active infection. However, our personal experience working with the parasite under similar conditions suggests it can achieve similar rates of infection regardless of the host being previously infected. Along with the results from the other groups, it seems likely that Met's infection rates observed in our experiment, and in the early exposure in particular, simply demonstrate the potential variation in the infectivity of this species. In such case, the positive contribution of Ht's vertical infection is by increasing the lower bar of Met's success if the conditions are suboptimal.

**Table S1. Infection rates of Met-exposed groups and their multiple comparisons.** In part A, treatments are classified by their exposure time and type (linked to Table 1 in the main text). Infection probability represents the proportion of infected individuals of a single group. In part B, comparisons are classified by their exposure time. The p-values are the result of Fisher's exact test and the effect sizes (ES) are the absolute value of the log of the odds-ratio estimated by the test. Single always means Ht absent, other types mean Ht present while vertical means Met exposure is sequential to vertical infection by Ht . Early and Late always refer to the time the animal was exposed to Met.

| **A.** | **Exposure time** | **Exposure type** | **Infection probability** |
|---|---|---|---|
| | **Early** | Single | 0.646 |
| | | Simultaneous | 0.48 |
| | | Sequential | 0.729 |
| | | Vertical | 0.957 |
| | **Late** | Single | 1 |
| | | Sequential | 0.98 |
| **B.** | **Exposure time** | **Comparison** | **p value**<br>**Effect size** |
| | **Early** | Single ~ Simultaneous | p = 0.13<br>ES = 0.674 |
| | | Single ~ Sequential | p = 0.51<br>ES = 0.386 |
| | | Single ~ Vertical | p < 0.001<br>ES = 2.489 |
| | | Simultaneous ~ Sequential | p = 0.021<br>ES = 1.059 |
| | | Simultaneous ~ Vertical | p < 0.001<br>ES = 3.316 |
| | | Sequential ~ Vertical | p = 0.007<br>ES = 2.103 |
| | **Late** | Single ~ Sequential | p = 1<br>ES = ∞ |

*Infection rate of Ht in groups nested within exposure times or within presence\absence of competitor*

**Table S2. Infection rates of Horizontal Ht-exposed groups and their multiple comparisons.** In part A, treatments are classified by their exposure time and type (linked to Table 1 in the main text). Infection probability represents the proportion of infected individuals of a single group. In part B, comparisons are classified by their exposure time. In part C, comparisons are classified both by exposure time and type. The p-values are the result of Fisher's exact test and the effect sizes (ES) are the absolute value of the log of the odds-ratio estimated by the test. Single always means Met absent, other types means Met present. Early and Late always refer to the time the animal was exposed to Ht.

| **A.** | **Exposure time** | **Exposure type** | **Infection probability** |
|---|---|---|---|
| | **Early** | Single | 0.49 |
| | | Simultaneous | 0.08 |
| | | Sequential | 0 |
| | **Late** | Single | 0.938 |
| | | Sequential | 0.292 |
| **B.** | **Exposure time** | **Comparison** | **p value**<br>**Effect size** |
| | **Early** | Single ~ Simultaneous | $p < 0.001$<br>ES = 0.093 |
| | | Single ~ Sequential | $p < 0.001$<br>ES = ∞ |
| | | Simultaneous ~ Sequential | $p = 0.118$<br>ES = ∞ |
| | **Late** | Single ~ Sequential | $p < 0.001$<br>ES = 3.567 |
| **C.** | **Met** | **Comparison** | **p value**<br>**Effect size** |
| | **Absent** | Early ~ Late | $p < 0.001$<br>ES = 2.741 |
| | **Present** | Simultaneous ~ Sequential (Early) | $p = 0.118$<br>ES = ∞ |
| | | Simultaneous ~ Sequential (Late) | $p = 0.013$<br>ES = 1.54 |
| | | Sequential (Early) ~ Sequential (Late) | $p < 0.001$<br>ES = ∞ |

*Spore production – additional results*

**Table S3. Additional effects on parasite spore production.** Summary of additional analysis of spore production. For each parasite, the analyses of two factors are given in two rows. The upper rows are the results for time of exposure and the lower rows for the non-infective presence of the competitor. Averages, standard errors and sample sizes are given for each level within a factor. Effect sizes were estimated by the normalized differences between levels. All p-values are derived from linear regression model, except for the p-value of Ht non-infective competitor that was calculated using Wilcoxon's test. The $p_{all}$ of Ht exposure time analysis represents the value of both the regression model and of all multiple comparisons. Effect sizes of this analysis are presented for each pair with level abbreviation as subscripts. Abbreviations: E= early, L= late, V= vertical, Abs= absence and Prs= presence.

| **Parasite** | **Average ± SE (n)** | **p-value**<br>**Effect size (ES)** |
|---|---|---|
| **Met** | E= $178.78x10^3 \pm 8.93x10^3$ (82)<br>L= $173.23x10^3 \pm 7.63x10^3$ (95) | $p = 0.47$<br>ES = 0.031 |
| | Abs= $181.31x10^3 \pm 8.93x10^3$ (75)<br>Prs= $171.75x10^3 \pm 5.36x10^3$ (102) | $p = 0.288$<br>ES = 0.053 |
| **Ht** | E= $7.83x10^3 \pm 3.49x10^3$ (24)<br>L= $271x10^3 \pm 38.64x10^3$ (46)<br>V= $861.53x10^3 \pm 36.69x10^3$ (49) | $p_{all} < 0.001$<br>$ES_{E\sim L} = 0.971$<br>$ES_{E\sim V} = 0.991$<br>$ES_{L\sim V} = 0.685$ |
| | Abs= $181.04x10^3 \pm 29.47x10^3$ (70)<br>Prs= $250.77x10^3 \pm 45.58x10^3$ (14) | $p = 0.13$<br>ES = 0.385 |

##### *Differences between Ht transmission modes in single infection*

The two transmission modes of Ht exhibited significant differences in their performance in and effects on their host. The vertical mode was significantly more virulent and produced more spores than the horizontal mode (Tables 3 and S2). While the horizontal mode had a very small effect on host reproduction and low spore production, the vertical mode reduced it dramatically and produced up to two orders of magnitude more spores. Vizoso and Ebert [1,2] suggested that there is a trade-off between the horizontal and vertical components of parasite transmission as well as different investment in within-host reproduction that depends on the transmission mode. Our results support this hypothesis and it is possible that each parasite transmission mode comes with its own reproductive strategy. When infecting horizontally, the parasite invests more resources in within-host expansion, taking over the reproductive system and infecting host offspring. The vertically-transmitted parasites are already well-established and thus can invest their resources in horizontal transmission, which is crucial for parasite expansion in the host population [3]. Interestingly, though producing different amounts of spores, the relative rate of their accumulation was similar in our model estimations (see below). This may further support the idea that the ability to exploit host resources is the same in both modes, but they are invested in different directions. Our findings are echoing those of Ben Ami et al. [4] and while genetic variation may produce variation in parasite virulence and its location on the horizontal-vertical tradeoff axis, we believe that the differences in reproductive strategy between Ht modes is a true characteristic of the parasite.

### Host survival analysis – additional information

#### *Calculations of effect sizes for Kaplan-Meir models*

Our first estimator of effect size in host survival models was the area under the curve (AUC) of the survival plot. It represents the cumulative survival probability of the group over time, and while being in strong alignment with the group's mean survival (Table 3 in the main text), it derives directly from the model. We calculated the AUC using the trapezoidal rule with a small adjustment. Since survival plots are comprised from rectangular bins (survival is constant between events), we computed a rectangular area rather than a trapezoidal one. Specifically, we took the events (in time) and probabilities vectors from the model and added the experiment's starting point (i.e., time=0, probability=1) to obtain a comparable framework. Then, we calculated the AUC as $\sum \Delta t_{i,i+1} * p_i$, where Δt is the duration from time i to time i+1 and p is the survival probability during this time. We used the normalized differences of AUC values as the effect size of the difference among categories for this measurement. We used the same outputs (events, probabilities and starting point) of the model to calculate host mortality rates. The regular (unweighted) mortality rate was calculated as the mean slope of the curve, i.e., $\frac{1}{n}\sum_{i=1}^{n}\frac{\Delta p_{i,i+1}}{\Delta t_{i,i+1}}$ where Δp and Δt are the difference between the value at time i+1 and I for probability and time, respectively, and n is the number of elements. The weighted mortality rate was calculated as the weighted mean slope of the curve, i.e., $\frac{\sum \Delta p_{i,i+1}}{\sum \Delta t_{i,i+1}}$ . The difference between these two rates is that the regular one is more sensitive to the number of events and represents better the intensity of mortality, while the weighted one is more sensitive to the length of events and represents differences that occur later on the time line, thereby creating a better predictor for future survival of individuals with extreme longevities. For example, survival probability can be estimated by $p_t$=mt + 1, where p is the survival probability at time t, m is the mortality rate and 1 is the survival probability at t=0. Therefore, the time it takes to reach survival probability of 0 will be t=-1/m. In addition, we used the rates ratio as the effect size of differences in mortality rate between groups. The regular rate of Met was -0.053, hence the time it takes to reach p=0 is -1/-0.053≈19, which is a close approximation for the time that passed from the day that first mortality occurred to the time that most individuals infected with Met died (Figure 4). The weighted rate of Met was -0.15, which gives a time estimate of 67 days, only a day shorter from the death of the last individual. Therefore, most individuals died fast within a short time span, while some managed to tolerate the parasite and significantly outlive their peers. In the control group, the regular rate resulted in time estimates of 63 days, or 83 days when measuring from the first observed death in the group. While this time period is close to the mean survival of the group (Table 3), half of the individuals lived longer than 80 days and about a quarter of them were alive at the end of the experiment (97 days). The weighted rate resulted in a time estimate of 125 days, which suggests that their longevity could span much beyond 80 days and allows predicting the max longevity of this group. Hence, the two estimates of mortality rate complement each other and provide a better understanding of the distribution of mortality at the group level.

##### *Results of the base model and follow-up comparisons*

Our first general Kaplan-Meir model with multiple paired comparisons demonstrated that all infection categories affected host survival ($p<0.001$ for all comparisons vs. control) and that Met and coinfection categories significantly from Ht ($p<0.001$ for both comparisons) but not from each other ($p= 0.325$). The effect size of Ht category on host survival (ES=0.165) was much smaller than Met and coinfection ($ES_{Met}$=0.639, $ES_{Coinfected}$= 0.652). Interestingly, the mortality rate of Ht was similar to the control category (ES=0.974), while the weighted rate showed a small difference (ES=1.207). This suggests that the effect of Ht starts long after the infection was initiated, or there exists a large variation in survival among Ht groups or both. To account for this potential variation, we further analyzed the Ht category by stratifying the survival data of animals infected with Ht, once by the age of exposure to the parasite and once by exposure to Met. Log-rank tests of each model demonstrated that exposure to Met did not affect the survival of Ht infected animals ($p=0.26$). However, host age at exposure significantly affected host survival of *Daphnia* infected with Ht ($p<0.001$ for the full model and each pairwise comparison). Analyzing the curves of the different times (vertical, early horizontal and late horizontal) and comparing them to the control group revealed large, exposure time dependent, variations in the effect size of this parasite on host survival (Table 4). Therefore, we split the Ht category in the final structure of the model to account for the different exposure times.

### Host reproduction – additional information

#### *The effect of parasites on host reproduction in the perspective of time*

To estimate the direct effect of the parasites on *Daphnia*'s clutch size, we had to validate that clutch size changes with host age and that infection by parasites can affect these changes. We calculated host age at every clutch they produced, which resulted in a repeated measure structure of individuals' reproduction. Since the data were very noisy, we flattened host age by assigning days to each week (e.g., ages of 0-6 days were assigned to week 1, 7-13 to week 2, and so on). We took the control category data and used a generalized mixed model (GLMM) with a Poisson distribution to test if there is an effect of host age in weeks on clutch size, while the individuals' identity was treated as a random effect. We estimated reproduction dynamics of each category by calculating the slope between consecutive weeks. We calculated the mean clutch size of each category for every week (except for week 1 where no reproduction occurred) and set the slopes to be the difference between the mean clutch sizes. We used a linear mixed model to test if the relative changes in clutch size were affected by the infection category, where week was used as a random effect (since the expected change is nested within the week). Following the conclusion of these models, we constructed a new model to test for the effect of parasites on host reproduction that considered the effect of host age. We used GLMM with Poison distribution with clutch size and infection category as the dependent and explanatory variables, respectively. Individuals' identity was used as a random effect to control for repeated measures and the week was used as another random effect to control for the effect of time.

To ensure that the estimate of the effect of the parasite was limited to specific time frames (i.e., consistent), we analyzed the AUC of the reproduction curves. We used the trapezoidal rule to calculate the AUC between each pair of consecutive weeks (from 2 to 14). Each AUC was calculated as $(S_t + S_{t+1})*\Delta t/2$, where S is the mean clutch size at time t and t+1 and $\Delta$t is time length between t and t+1. Since the time step was constant and equal to 1 then $AUC_{t,\,t+1}= 0.5*(S_t + S_{t+1})$. We calculated the effect size of each AUC from each infection category against its respective AUC from the control curve as the normalized AUC difference. To determine if the resulting effect sizes are of statistical significance, we used a bootstrap-style simulation to generate a domain of possible effect sizes of our control group against a virtual equivalent. We started by extracting the max observed clutch size of each week and used it to generate an array of possible clutch size for each week spanning from 0 to the respective max size. For week 3, we corrected the max value (27) to be the second highest value (11) since the max value was an extreme size outlier (mean ± SD = 5.739 ± 3.6). We simulated a virtual reproduction curve by computing, for each week, the mean clutch size of 50 "individuals" (the starting n of a group in our experimental design) whose clutch size was randomly drawn from the possible values array of the respective week. We calculated AUCs of the virtual curve, computed their effect size from the observed and added the results to the effect sizes domain. We repeated this simulation for 1000 iterations to end up with an effect sizes domain with 12000 values. We estimated the p-values of our observed effect sizes as the number of values that are equal or larger/smaller (for positive/negative effect sizes, respectively) in the effect sizes domain divided by the total number its values.

*Results of the base model and final infection category's structure*

The results of our base linear regression model (infection categories: control, Ht, Met and coinfection) demonstrated that the infection categories had a significant effect on host clutch size ($p<0.001$) and all groups differed from each other in the post-hoc multiple comparisons ($p<0.001$ for all comparisons). For the categories Met and Ht that were pooled from several treatment groups, we tested if factors that affected parasite performance in previous analyses resulted in variation in parasite effect on the host's mean clutch size. We used a linear model to test if Met exposure time affected infected hosts clutch size and found no effect ($p=0.791$). Similarly, we used a linear model to test if exposure time to Ht or the presence of Met at exposure affected Ht infected hosts' average clutch size. While the presence of Met had no effect on host mean clutch size ($p=0.583$), exposure time had a significant effect on the clutch size of Ht infected hosts ($p<0.01$) and all multiple comparisons were significant ($p<0.001$ for vertical vs. both early and late horizontal, $p=0.008$ for early vs. late). However, the size difference between the two horizontal exposures was 1.5 offspring with a small effect size (ES=0.118). Hence in the structure of the final model, we split the Ht category to vertical and horizontal and used the new category structure (control, vertical Ht, horizontal Ht, Met and coinfected) in all subsequent models.

*The dynamics of clutch size and host age*

Clutch size of control individuals changed significantly with host age (Wald=1400.6, $p<0.01$). The size dynamic was characterized by small clutches at the onset of reproduction, a large prime peak when *Daphnia* are fully matured and smaller secondary peaks at older age (Figure 4). Each peak was followed by an exhaustion phase of reduction in clutch size (Figure 4). An increase in clutch size during the last week of the experiment may suggest another cycle of peak and exhaustion, but we cannot be sure. These dynamics were not affected by the different infection categories ($p>0.8$ for all multiple comparisons).

### Effects of exposure to parasites that did not result in an infection

Seventy-one individuals from all exposure categories (besides those involving vertical transmission by Ht) did not develop an infection. Therefore, we decided to test if mere exposure to parasites had any effect on host fitness (The effect of this scenario on parasite fitness is discussed in the main text). We used similar methodology to that described in the main text to test if these "exposed but uninfected" individuals differed from the control group in the following traits: host survival (Kaplan-Meir), onset of reproduction (GLM with gamma distribution), average length between clutch interval (LM) and average clutch size (LM). Since exposure did not affect host survival (see below), there was no need to adjust the clutch size analysis to control for the effect of time. We tested each parameter twice, once by comparing the control to the pool of all exposed but not infected individuals, and once by comparing the control to each type of exposure (only to Met, only to Ht or to both). None of the models (with pooled or detailed exposure types) yielded significant results. Therefore, we concluded that exposure to parasites without becoming infected did not affect the fitness of *Daphnia* in our experiment. The results of all models are summarized in Table S4.

**Table S4. Effects of exposure without infection.** Results of all tested models tested of the effects of parasite exposure that did not result in an infection on host fitness. Averages, standard errors and sample size are given for every group tested against the control group. The p-values and effect sizes (ES) are given per model. Effect sizes are estimated by the normalized differences of the groups' averages from the control group (see caption of Table 3). The effect sizes of the detailed exposures models are the mean of the absolute value of the effect sizes of each group against the control.

| Parameter | Group | Mean ± SE (n) | p-value<br>Effect size |
|---|---|---|---|
| **Mean survival** | Negatives | 71.099 ± 2.893 (71) | p = 0.99<br>ES = 0.053 |
| | Exposed to Met | 79.765 ± 5.781 (17) | p = 0.49<br>ES = 0.08 |
| | Exposed to Ht | 68 ± 4.216 (28) | |
| | Exposed to both | 68.769 ± 5.171 (26) | |
| **Age of first event** | Negatives | 16.225 ± 0.284 (71) | p = 0.799<br>ES = 0.007 |
| | Exposed to Met | 17.412 ± 0.762 (17) | p = 0.112<br>ES = 0.042 |
| | Exposed to Ht | 15.679 ± 0.463 (28) | |
| | Exposed to both | 16.038 ± 0.269 (26) | |
| **Average interval between events** | Negatives | 4.029 ± 0.09 (70) | p = 0.555<br>ES = 0.018 |
| | Exposed to Met | 4.311 ± 0.23 (17) | p = 0.083<br>ES = 0.045 |
| | Exposed to Ht | 4.07 ± 0.124 (28) | |
| | Exposed to both | 3.791 ± 0.129 (25) | |
| **Average clutch size** | Negatives | 12.753 ± 0.346 (71) | p = 0.374<br>ES = 0.034 |
| | Exposed to Met | 12.148 ± 0.775 (17) | p = 0.191<br>ES = 0.057 |
| | Exposed to Ht | 13.52 ± 0.433 (28) | |
| | Exposed to both | 12.324 ± 0.634 (26) | |
